# Cracked actin filaments as mechanosensitive receptors

**DOI:** 10.1101/2023.06.26.546553

**Authors:** Vilmos Zsolnay, Margaret L. Gardel, David R. Kovar, Gregory A. Voth

## Abstract

Actin filament networks are exposed to mechanical stimuli, but the effect of strain on actin filament structure has not been well-established in molecular detail. This is a critical gap in understanding because the activity of a variety of actin-binding proteins have recently been determined to be altered by actin filament strain. We therefore used all-atom molecular dynamics simulations to apply tensile strains to actin filaments and find that changes in actin subunit organization are minimal in mechanically strained, but intact, actin filaments. However, a conformational change disrupts the critical D-loop to W-loop connection between longitudinal neighboring subunits, which leads to a metastable cracked conformation of the actin filament, whereby one protofilament is broken prior to filament severing. We propose that the metastable crack presents a force-activated binding site for actin regulatory factors that specifically associate with strained actin filaments. Through protein-protein docking simulations, we find that 43 evolutionarily-diverse members of the dual zinc finger containing LIM domain family, which localize to mechanically strained actin filaments, recognize two binding sites exposed at the cracked interface. Furthermore, through its interactions with the crack, LIM domains increase the length of time damaged filaments remain stable. Our findings propose a new molecular model for mechanosensitive binding to actin filaments.

**SIGNIFICANCE STATEMENT:** Cells continually experience mechanical strain, which has been observed to alter the interactions between actin filaments and mechanosensitive actin-binding proteins in recent experimental studies. However, the structural basis of this mechanosensitivity is not well understood. We used molecular dynamics and protein-protein docking simulations to investigate how tension alters the actin filament binding surface and interactions with associated proteins. We identified a novel metastable cracked conformation of the actin filament, whereby one protofilament breaks before the other, presenting a unique strain-induced binding surface. Mechanosensitive LIM domain actin-binding proteins can then preferentially bind the cracked interface, and this association stabilizes damaged actin filaments.

## INTRODUCTION

Eukaryotic cells utilize the actin cytoskeleton to move, change and maintain shape, divide, and transport cargo. These processes subject the underlying actin filament (F-actin) cytoskeleton to a wide range of mechanical forces including tension, bending, compression, and twisting (1, 2). Cells utilize mechanotransduction pathways to exploit mechanical cues to modify biochemical activity, which enables cells to alter cell physiology, transcription, and actin dynamics (1-3). For example, the actin-binding proteins (ABPs) Arp2/3 complex (4, 5), α-catenin (6), and in some studies, cofilin (7, 8), have been reported to exhibit altered binding affinities for actin filaments subjected to mechanical strain. These altered ABP interactions have important downstream implications for network architecture, stiffness, and disassembly (1, 2).

At the molecular level, actin filaments are composed of two protofilament strands that form a double helix (Fig. 1A). Individual actin subunits within the filament form inter-strand (lateral) or intra-strand (longitudinal) interactions with neighboring subunits. A number of cryo-electron microscopy (cryo-EM) structures have recently been reported of actin filaments up to a resolution of 2.2 Å (9-12). These structures have facilitated an unprecedented understanding of the conformation of actin filaments under unstrained conditions. However, much less is known about how the conformation of the actin filament changes under strain. A pioneering cryo-EM study recently reconstructed bent actin filaments at a resolution of ∼3.6 Å (12). These structures revealed deformations to the actin filament lattice consistent with the twist-bend coupling predicted by theoretical studies (13) and did not find that the filament surface charge or hydrophobicity changes in a meaningful way. As such, the molecular mechanism by which binding of mechanosenstive ABPs is activated by strain remains an open question.

**Figure 1.**
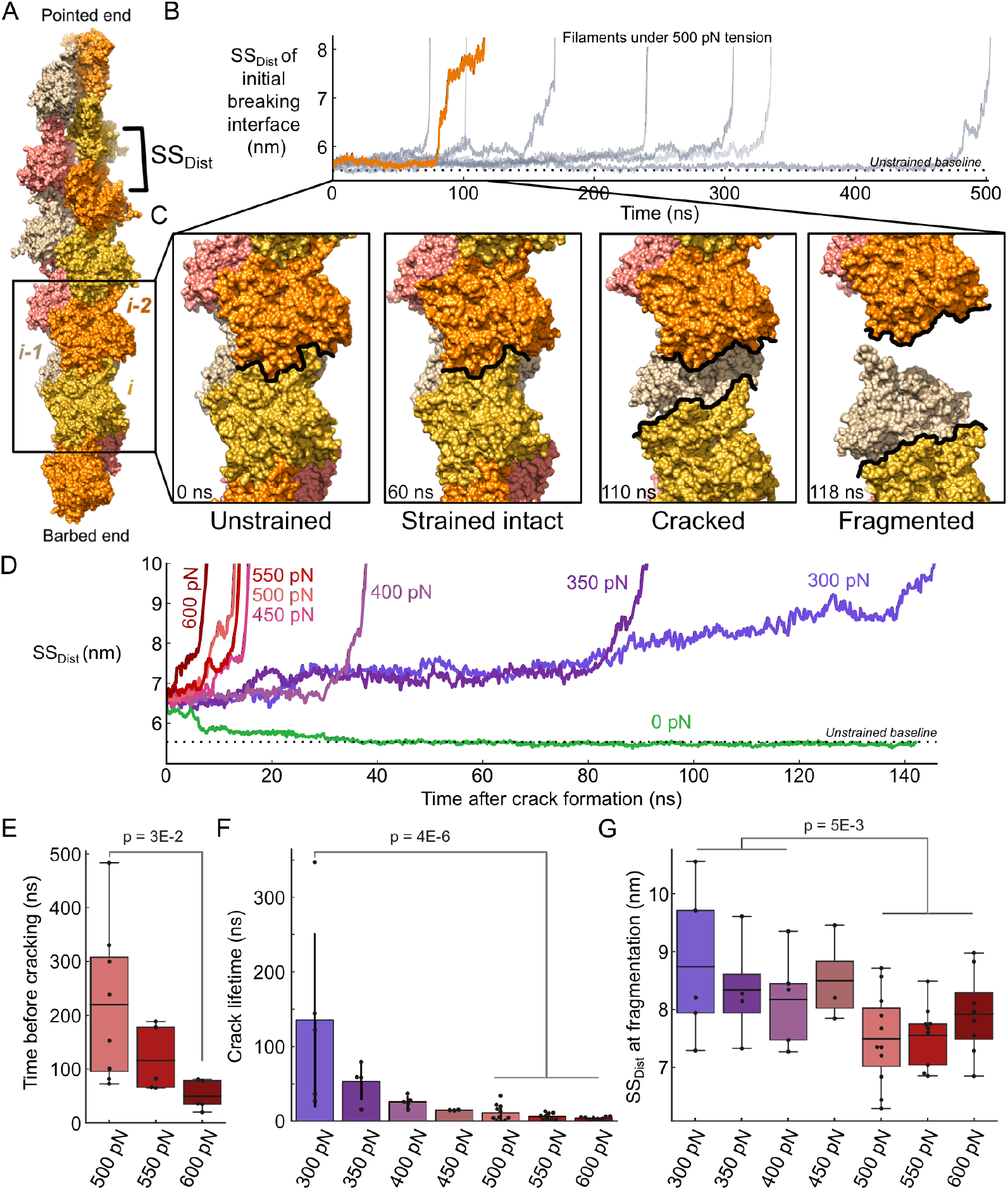
Actin filaments enter a metastable ‘cracked’ state prior to fragmentation. (A) Space-filling model of the initial structure of an ADP-actin 13-mer. The two strands of the protofilament are colored in alternating orange and yellow, and pink and tan, respectively. The distance between longitudinal neighboring subunits, subunit-subunit distance (ss_dist_), is labeled. (B) Time courses of the ss_dist_ at the initial breaking interface in eight independent simulations run at 500 pN tension (Table S1). (C) Space-filling model snapshots of a representative simulation (orange time course in A) depicting the initial breaking interface in unstrained, strained-intact, cracked, and fragmented conformations. This process is shown in Movie S1. (D) Time courses of the ss_dist_ of cracked interfaces run under different magnitudes of tension illustrate a tension-dependent lifetime of cracks. In the absence of tension, cracked interfaces spontaneously return to unstrained baseline values (green). (E) Box plots of the time before the initial interface cracks reveal an inverse relationship between the applied force and the time the filament remains intact. n_500_ = 8 simulations, n_550_ = 5, n_600_ = 5. (F) Bar plots of the time spent in the cracked state reveal that the crack lifetime is inversely correlated with the magnitude of the tension applied. Error bars represent standard deviation. The p-value is calculated between 300 pN and cases ≥ 500 pN. n_300_ = 5 simulations, n_350_ = 4, n_400_ = 5, n_450_ = 3, n_500_ = 11, n_550_ = 9, n_600_ = 8. (G) Box plots of the ss_dist_ at the time of fragmentation show that at lower tensions filaments fragment with larger separations at the cracked interface. The p-value is calculated between cases ≤ 400 pN and cases ≥ 500 pN. n_300_ = 5 simulations, n_350_ = 4, n_400_ = 5, n_450_ = 3, n_500_ = 11, n_550_ = 9, n_600_ = 8.

For example, recent studies have identified that proteins in the LIM (Lin11, Isl-1, and Mec-3) domain family localize to actin filaments under mechanical strain (14, 15). Structurally, LIM domains are ∼ 60 amino acid sequences containing two zinc finger motifs, and many LIM proteins contain multiple LIM domains in tandem separated by flexible linkers of 7-8 amino acids (15). These studies observed that the LIM-containing region (LCR) of these proteins alone is sufficient for force-activated binding to F-actin *in vivo* and *in vitro*, but that multiple tandem LIM domains with characteristic spacings are required for mechanosensitivity (14, 15). A subsequent study reported that a single LIM domain of testin is mechanosensitive on its own (16), but presently this is the only known exception. Despite a growing body of experimental literature investigating the behavior of a diverse set of LIM domain proteins, a mechanistic understanding of how LIM recognizes strained F-actin remains unknown.

Here we use all-atom molecular dynamics (MD) simulations to investigate the structure and dynamics of actin filaments under applied tension. We find that Y169 of one subunit flips away from the D-loop of its longitudinal neighboring subunit, which leads to the breaking of a single protofilament prior to filament fragmentation. This metastable cracked interface alters the filament binding surface, which mechanosensitive LIM domains preferentially associate with and their presence stabilizes damaged filaments. Our results elucidate key steps of the filament fragmentation pathway, lead to natural explanations for why multiple tandem LIM domains increase mechanosensitive binding, and provide a new molecular model for mechanosensitive binding to F-actin generally.

## RESULTS

### Actin filament fragmentation involves a metastable ‘cracked’ state

To determine the molecular rearrangements induced by mechanical strains, we built an MD system consisting of an actin filament composed of 13 actin subunits (see Methods). The centers of mass of the three terminal subunits at the barbed and pointed ends were pulled away from each other with a specified applied force. This resulted in 9 longitudinal interfaces (between subunits i and i-2) subjected to mechanical tension (Fig. 1A). In total, we performed 55 simulations of the actin 13-mer filament at varying tensions (Table S1).

In all-atom MD simulations of actin filaments under applied tension, strains of 500 pN and above cause filaments to fragment within timescales accessible by large all-atom MD (∼1 µs), whereas tensions 400 pN and below do not result in breaks on this timescale (Table S1). Interestingly, filament fragmentation does not occur as a single discrete event. Instead, longitudinal connections between subunits on each protofilament break separately. At all longitudinal interfaces, including the first interface to break, the subunit-subunit distance (ss_dist_) initially remains close to the unstrained baseline value of ∼ 5.53 ± 0.02 nm (Fig. 1B) and maintains the typical longitudinal connections of unstrained filaments (Fig. 1C *left* and *middle left, Unstrained* and *Strained intact*). After a period where all strained interfaces remain intact, one longitudinal interface loses contact and separates while interfaces on the other protofilament maintain their connections, which results in the formation of a metastable ‘crack’ (Fig. 1C *middle right, Cracked*, Movie S1). Under continued tension, one of the laterally neighboring interfaces eventually loses contact and the filament fully fragments (Fig. 1B, C *right, Fragmented*).

The timing of the steps along the fragmentation pathway is modulated by tension. Increasing the strain applied to the filament after the crack forms decreases the amount of time before the filament fragments (Fig. 1D, F). Conversely, cracks spontaneously reform lost contacts in the absence of applied tension (Fig. 1D, green line). Additionally, the time preceding the initial break (i.e., the crack) inversely correlates with strain (Fig. 1E). The ss_dist_ of the cracked interface at fragmentation trends with tension, whereby cracks under higher tensions result in filament fragmentation at lower separations (Fig. 1G).

### Actin filaments break at interfaces where Y169 flips away from the D-loop

In unstrained actin filaments, the D-loop of subunit i completely envelopes Y169 of the neighboring subunit i-2 (9, 10). This connection provides a substantial fraction of the buried surface area between subunits (9) and is the primary connection between the terminal barbed end subunit and the rest of the filament (17). Under tension, the initial breaking interface is characterized by Y169 of subunit i-2 flipping away from the D-loop of subunit i (Fig. 2A, Movie S2). The flip of Y169 increases the distance between Y169 and the D-loop from ∼4 to >7 Å (Fig. 2B *top*) and precludes the typical enveloping interaction. In turn, the number of contacts Y169 forms with residues in subunit i decrease from ∼10 to ∼4 (Fig. 2B *middle top*). As the important Y169 to D-loop connection erodes, the ss_dist_ of the interface increases (Fig. 2B *middle bottom*) in step with a marked decrease of the overall number of contacts between subunits i and i-2 (Fig. 2B *bottom*).

**Figure 2.**
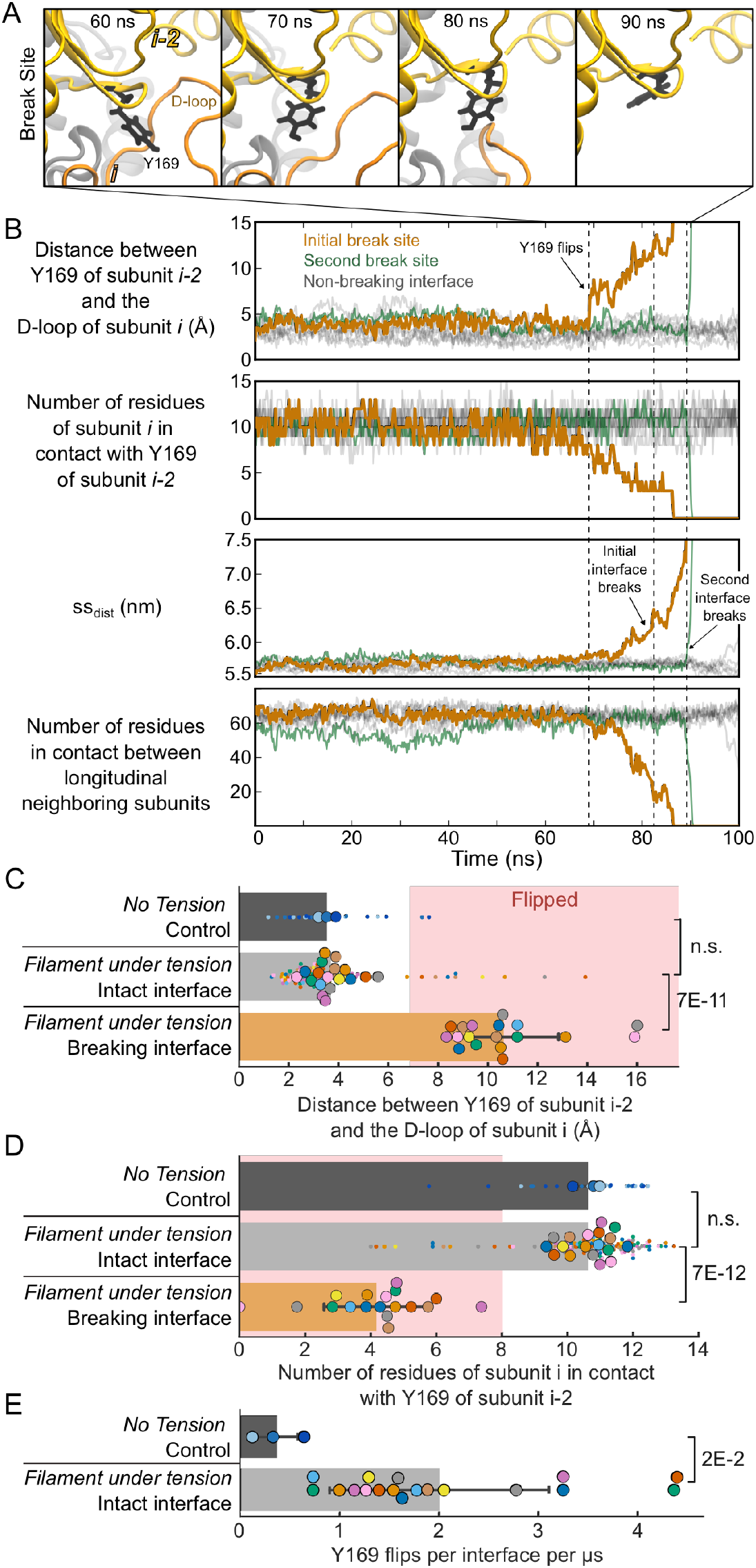
Actin filaments break at interfaces where Y169 flips away from the D-loop of the longitudinally neighboring subunit. (A) Snapshots of a representative longitudinal interface breaking under 550 pN of tension. The break (90 ns) is preceded by Y169 of subunit i-2 (black) flipping away (between 60 and 70 ns) from its connection with the D-loop of subunit i (orange). This process is illustrated in Movie S2. (B) Time courses of key metrics tracking a longitudinal interface breaking under 550 pN of tension from a representative simulation. The first and second interfaces to break are shown in orange and green, respectively, and non-breaking interfaces are shown in gray. Dashed lines mark the annotated transitions. (Top) The distance between the CZ atom of Y169 of subunit i-2 and the center of mass of Cα of residues 41, 42, 48, 49 of the D-loop of subunit i at each interface reveals a characteristic flip of Y169 at the initial break site. (Middle Top) After flipping, the number of residues of subunit i in contact with Y169 of subunit i-2 decreases. (Middle Bottom and Bottom) The distance between the center of masses of subunits i and i-2, ss_dist_ (Middle Bottom), at the initial break site increases in step with a corresponding decrease in the total number of contacts (Bottom), leading to complete fragmentation. Contacts are calculated using a 5 Å cutoff. (C and D) Swarm plots of the distance between the CZ atom of Y169 of subunit i-2 and the center-of-mass of the D-loop of subunit i (C), as well as the number of inter-subunit contacts with Y169 (D). Intact interfaces with and without tension (dark and light gray bars, respectively) both generally maintain unflipped Y169 residues with a small population of flipped residues. In contrast, Y169 is flipped at all breaking interfaces, which results in reduced contacts (orange bars). Values are averages of the final 2% of frames preceding the initial break. Small dots represent individual interfaces, while large dots represent the mean across all interfaces in a given simulation; color corresponds to replicates. Error bars represent standard deviation; p-values are displayed on the right. n_control_ = 3 simulations (27 interfaces), n_intact_ = 18 simulations (144 interfaces), n_breaking_ = 18 simulations (18 interfaces). (E) Swarm plots of the rate of Y169 flipping reveal that tension increases the rate of flipping events. Error bars represent standard deviation; the p-value is displayed on the right. n_control_ = 3 simulations (27 interfaces), n_intact_ = 18 simulations (162 interfaces).

Although the average interface maintains the low ∼ 4 Å Y169 to D-loop distance observed in cryo-EM structures (9-12), there is a small subpopulation of interfaces where Y169 is in the flipped orientation (Fig. 2C, *dark gray bar and small dots*). Interfaces under tension, but not at the breaking interface, show a similar distribution wherein most Y169 orientations are not flipped (Fig. 2C, *light gray bar and small dots*). However, at interfaces that break, Y169 is always in the flipped state at the time the break occurs (Fig. 2C, *orange bar*, 18 out of 18 independent runs). Intuitively, these trends are reflected in the number of connections Y169 of subunit i-2 forms with subunit i (Fig. 2D). Although Y169 flipping is possible in unstrained simulations, the rate of flipping increases ∼5 fold under tension (Fig. 2E). Flips of Y169 only correlate with broken longitudinal interfaces under applied tension and do not lead to cracks in unstrained simulations.

### The cracked interface presents a unique binding surface

When averaged over 3 simulations, unstrained filaments maintain an ss_dist_ of 5.53 ± 0.02 nm (Fig. 3A), and over a range of tensions strained-intact filaments show only a slight increase. For example, at 200 and 600 pN the ss_dist_ of intact interfaces is 5.57 ± 0.01 nm and 5.71 ± 0.03 nm, respectively (Fig. 3A). In contrast, when averaged over 45 simulations, cracked interfaces maintain much larger ss_dist_ values of 6.94 ± 0.37 nm (Fig. 3A).

**Figure 3.**
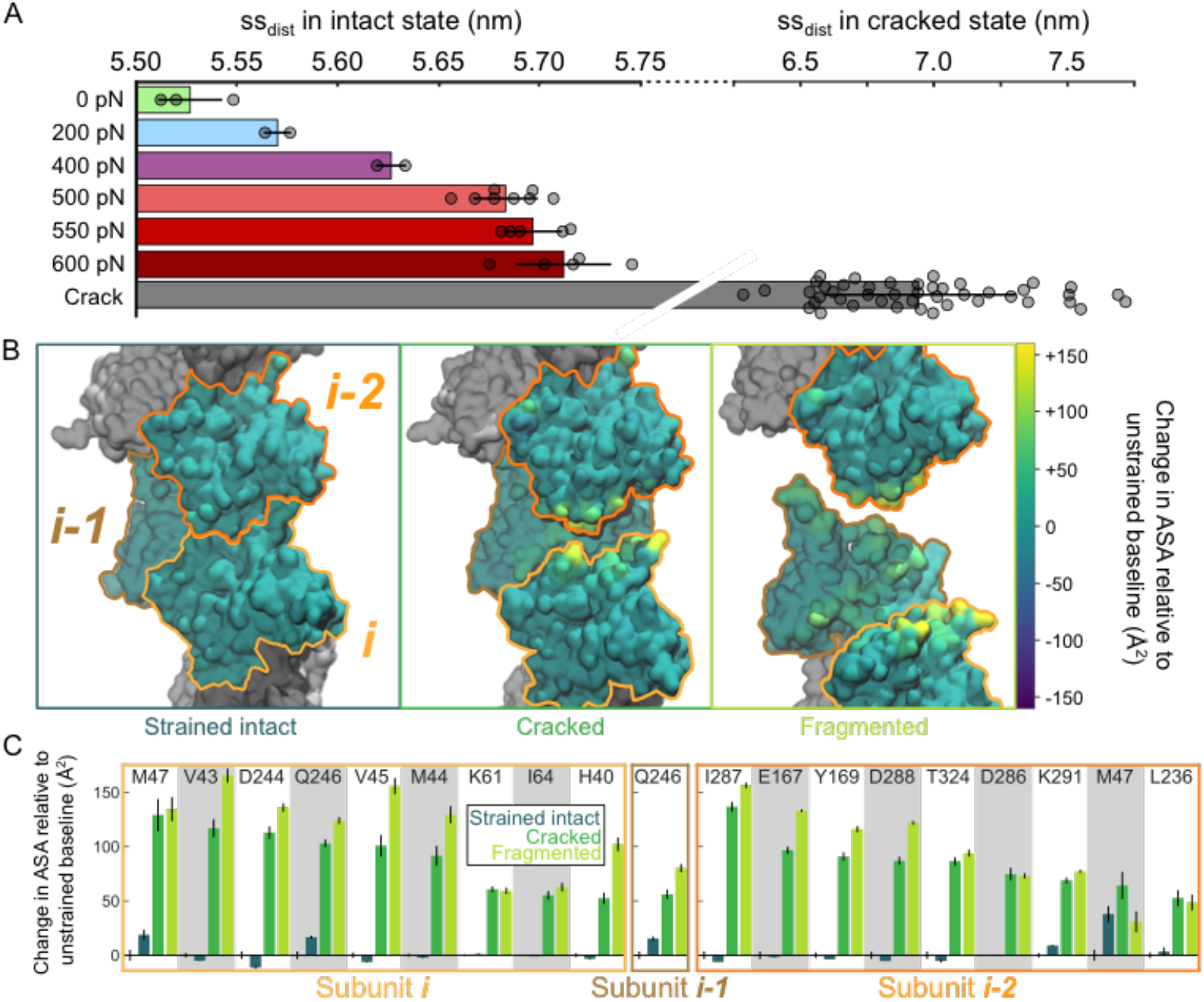
Surfaces of strained-intact filaments differ minimally from unstrained filaments, whereas cracks present a unique strain-induced binding surface. (A) Bar plots of the ss_dist_ reveal a consistent increase in the average distance between longitudinal neighboring subunits in intact filaments under tension, up to ∼0.2 nm. The ss_dist_ of cracked interfaces are shown for comparison; Note the broken axis. Error bars represent standard deviations. n_0_ = 3 simulations, n_200_ = 2, n_400_ = 2, n_500_ = 8, n_550_ = 5, n_600_ = 5, n_crack_ = 45. (B) Space-filling models of representative filament interfaces in the strained intact, cracked, and fragmented states with surfaces colored according to the change in accessible surface area (ASA) of residues relative to unstrained baseline interfaces. Strained-intact interfaces do not show major differences relative to baseline, whereas cracked interfaces expose many typically-buried residues, some of which are comparable to values of fragmented interfaces. Only changes in ASA that are significantly different from unstrained (p < 0.05) are included. n_intact_ = 22 simulations, n_cracked_ = 45, n_fragmented_ = 45. (C) Bar plots of the average change in ASA of specific residues (from subunits i, i-1 and i-2 as indicated) that show the greatest increase relative to subunits within unstrained filaments. Each residue is separated by alternating shaded regions and contains four bars corresponding to unstrained, strained intact, cracked, and fragmented interfaces. Unstrained is the baseline case and therefore has a mean of 0. The probe radius is 4 Å and residues are included if either the strained intact or cracked states differ from unstrained by more than 50 Å^2^; All residues are shown in Figs. S3, S4, and S5. Error bars represent standard error. n_intact_ = 22 simulations, n_cracked_ = 45, n_fragmented_ = 45.

Because cracked and strained-intact interfaces are both mechanically-induced states of the actin filament with the potential to present cryptic binding sites for mechanosensitive actin binding proteins, we evaluated how the filament surface in these states differed from unstrained filaments. In particular, we considered whether the slight increases observed in the ss_dist_ of strained-intact filaments expose residues to the filament surface that are buried in unstrained filaments. To quantify these changes, we performed accessible surface area (ASA) calculations of each amino acid at a probe radius of 4 Å. This revealed that the increased separation between longitudinal neighboring subunits in strained-intact filaments does not translate into large increases of the accessibility of individual amino acids (Fig. 3B *left* and S2). In contrast, cracked interfaces show meaningful increases in the accessibility of many typically-buried residues (Fig. 3B *middle* and S3), some of which become exposed to levels comparable to completely fragmented interfaces (Fig. 3B *right* and S4). Many of the largest gains are made by hydrophobic residues in the central D-loop of subunit i, V43, V45, M44, and M47 (Fig. 3C), which typically contact Y169 of subunit i-2 at intact interfaces. Summing over all residues, cracked interfaces have on average more than 11 times the increase in ASA of strained-intact interfaces, and over half the increase in ASA of fragmented interfaces (Figs. S2, S3, and S4).

### LIM domains bind cracked interfaces and stabilize filaments under tension

Given that several ABPs have been observed to have increased binding activity to strained actin filaments *in vivo* and *in vitro* and cracks present a highly unique filament surface (Fig. 3C), we tested whether various ABPs associate preferentially with cracked interfaces by performing protein-protein docking simulations using ClusPro2.0 (see Methods) (18). We used a representative cracked 13-mer filament structure that emerged from MD simulations as the receptor and individual ABP structures as the ligand molecule. A number of ABPs bind to actin in known ways without reported preferences for strained actin filaments. For instance, capping protein, profilin, and actin monomers preferentially bind to the barbed end of the actin filament in docking simulations, as expected, with little to no association with the cracked interface (Fig. 4B, S1A). In contrast, the structures of 43 evolutionarily diverse zinc-finger containing LIM domains predicted using Alphafold2 (see Methods) (19) overwhelmingly bind the cracked interface over other regions of the actin filament receptor, such as the filament ends and non-cracked interfaces (Fig. 4B and S1). This is particularly striking given that, in addition to one cracked interface, the actin 13-mer receptor contained ten strained-intact interfaces that were available for binding, which served as an internal control for each simulation.

**Figure 4.**
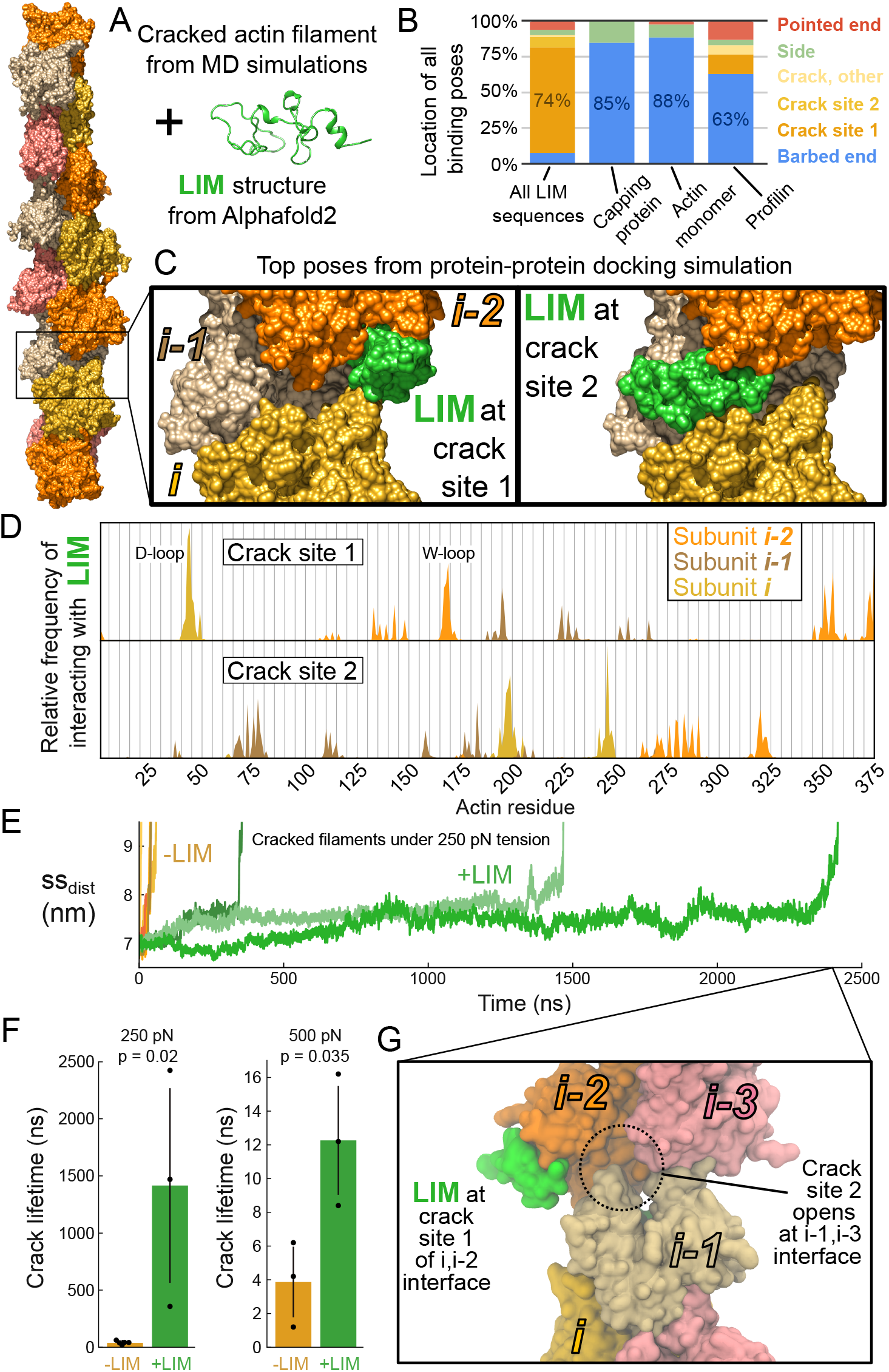
LIM domains bind cracked interfaces at two sites and stabilize filaments. (A) Left, Space-filling model of an actin 13-mer filament with a single cracked interface (boxed region). Right, Ribbon diagram of the first LIM domain of testin predicted by Alphafold2. The cracked filament and LIM domains were input into protein docking simulations as the receptor and ligand, respectively. (B) Location of all docking poses on a cracked actin 13-mer receptor for the indicated ligands. All 43 LIM domain sequences are shown at left as an aggregated dataset. The most likely binding location for each ligand is labeled as a percentage on the bar. Note that LIM domains have a unique preference for the cracked interface. Additional ligands are shown in Fig. S1A. (C) Space-filling models of representative output poses from docking simulations of the first LIM domain of testin bound to crack site 1 and crack site 2. Top poses for all 43 LIM domains bind the cracked interface rather than all other regions of the filament surface. (D) Histogram of the frequency that each actin subunit (i, i-1 and i-2 as indicated) residue is in contact (5 Å cutoff) with a LIM domain in docked structures at crack site 1 (top) and crack site 2 (bottom). Data is aggregated across all 43 LIM domains tested. (E) Time courses of the ss_dist_ of MD simulations of a cracked actin 7-mer with (green) and without (orange) the first LIM domain of testin bound to both crack sites under 250 pN of tension, revealing that damaged filaments are more stable in the presence of LIM domains. (F) Bar graph of the crack lifetime at two tensions show that the first LIM domain of testin stabilizes damaged actin filaments. Error bars represent standard deviation. n_250,–LIM_ = 5 simulations, n_250,+LIM_ = 3, n_500,–LIM_ = 3, n_500,–LIM_ = 3. (G) Snapshot of a representative actin filament with the first LIM domain of testin bound to the initial cracked interface reveals that crack site 2 of the interface on the opposing strand becomes exposed prior to fragmentation. This process is illustrated in Movie S3.

Individual LIM domains are predicted to bind the cracked interface in two locations (Fig. 4C). The vast majority of binding poses are located between subdomain 2 of subunit i and subdomains 1 and 3 of subunit i-2 (Fig. 4C), accounting for 93% of accepted poses across the 43 LIM domain sequences (Fig. S1C). In this position, LIM bridges the broken connection between the D-loop and the W-loop containing Y169 and makes contact with all three subunits at the cracked interface (Fig. 4D). In the second most common binding location, crack site 2, the LIM domain contacts subdomain 4 of subunit i and subdomain 3 of subunit i-2 simultaneously (Fig. 4C and D). Many of the key actin residues that LIM interacts with at the two crack sites (Figs. S5 and S6) only become exposed once cracks form (Fig. S3).

Intriguingly, a LIM domain bound to crack site 1 does not clash with a LIM domain bound to crack site 2, suggesting two LIM domains can bind a single cracked interface. This is particularly interesting because multiple LIM domains in tandem were observed to be an important driver of mechanosensitivity *in vivo* (14, 15). Having multiple, non-mutually exclusive LIM binding sites at the crack suggests a structural basis for LIM domain’s apparent avidity-driven mechanosensitive response.

We sought to test the idea that the presence of LIM at the cracked interface stabilizes the filament by itself. To that end, we built an MD system consisting of a cracked actin 7-mer with the first LIM domain of testin bound to each of crack sites 1 and 2, given that recent experiments reported that the first LIM domain of testin is mechanosensitive as an individual (i.e. not tandem) LIM domain. At both 250 pN and 500 pN tensions (14 simulations total), the presence of the two testin LIM domains significantly prolongs the period of time before the filament fragments (Fig. 4E and F). This is especially pronounced at 250 pN, where the presence of LIM domains at the cracked interface increases the time before fragmentation over 37-fold from 38 ± 16 ns to 1416 ± 841 ns (Fig. 4E and F). Interestingly, immediately before fragmentation, crack site 2 of the opposite protofilament became exposed (Fig. 4G, Movie S3). This suggests the possibility that after a crack has formed and been stabilized by LIM domains on one protofilament, continued tension can expose additional crack sites on the laterally neighboring interfaces available for further LIM binding.

## DISCUSSION

### Methodological considerations

We set out to observe the fragmentation of actin filaments on MD timescales, typically limited from hundreds of nanoseconds to microseconds for large, multiprotein systems. Therefore, we applied tensions on the order of hundreds of piconewtons, which is higher than most physiological tensions. However, several lines of evidence suggest that our major conclusions are relevant at lower forces. First, Y169 flipping is observed in control simulations in which no tension is applied (Fig. 2C and D). Tension increases the rate at which Y169 flips and results in the interface breaking, but is not required for flipping to occur (Fig. 2). Second, although we had not predicted the metastable cracked state before performing these simulations, it is intuitive in hindsight and should be relevant across all tensions. Actin is a double helix composed of two protofilament strands. As long as both protofilaments do not break at neighboring interfaces at the exact same time, cracks will form. The physiological relevance of the intermediate cracked state depends on how long cracks remain metastable, and it is clear from our data that cracks last longer, and so would be even more relevant, at lower tensions. Additionally, cracks in actin filaments are reminiscent of the physiologically relevant damaged states reported for other biological polymers such as microtubules and DNA (20, 21).

However, it is possible that the relatively high tensions that we applied here obscure the existence of additional steps on the fragmentation pathway (i.e., these steps did not have enough time to become noticeable in the MD runs). With the lowest tension in which we observed fragmentation, 250 pN, crack site 2 became exposed before crack site 1. It seems plausible that at lower tensions, crack site 2 consistently becomes exposed first, as the exposure of crack site 1 is limited by the flip of Y169. Importantly, we applied strain using constant force, which is an important choice for this and any similar study. Constant velocity pulling schemes will not reveal metastable states, as force is increased on-the-fly to push past such metastable states.

Additionally, we applied tensile strains. We suggest that our findings under these conditions may be relevant to other modes of strain, which at the scale of an individual interface likely appear very similar to tension (e.g., an out-of-equilibrium separation between neighboring subunits). In support of this, mesoscale modeling of actin filaments indicated that strain is concentrated under a variety of modes of applied force (e.g., bending, compression, tension, twisting) (22). We propose localized strain under other types of mechanical strain also leads to a similar metastable cracking behavior as observed in our all-atom MD simulations. However, further studies are required to investigate differences that may arise from different modes of applying force.

Lastly, docking simulations have important limitations. In particular, molecules are treated as rigid bodies, so flexible linkers or conformational changes upon binding are not accurately reflected (23). For this reason, we used the structured individual LIM domains instead of tandem LIM domains separated by flexible linkers as the ligands for our docking simulations, and focused on conclusions that were common among the 43 LIM domain sequences we tested.

### The connection between Y169 and the D-loop stabilizes actin filaments

We found that flipping of Y169 away from the neighboring subunit’s D-loop is a necessary step for longitudinal interfaces to break under strain (Fig. 2, 5A). This result adds to the evidence that the connection between Y169 of subunit i-2 and the D-loop of subunit i is critical for maintaining the stability of actin filaments (9, 17). In addition to residue 169 being highly conserved as either a tyrosine or phenylalanine in WT actin sequences, previous studies have demonstrated that the D-loop’s connection with Y169 buries a large fraction of the surface area between subunits within the polymer (9) and alone can support the terminal barbed end subunit’s attachment to the filament (17).

**Figure 5.**
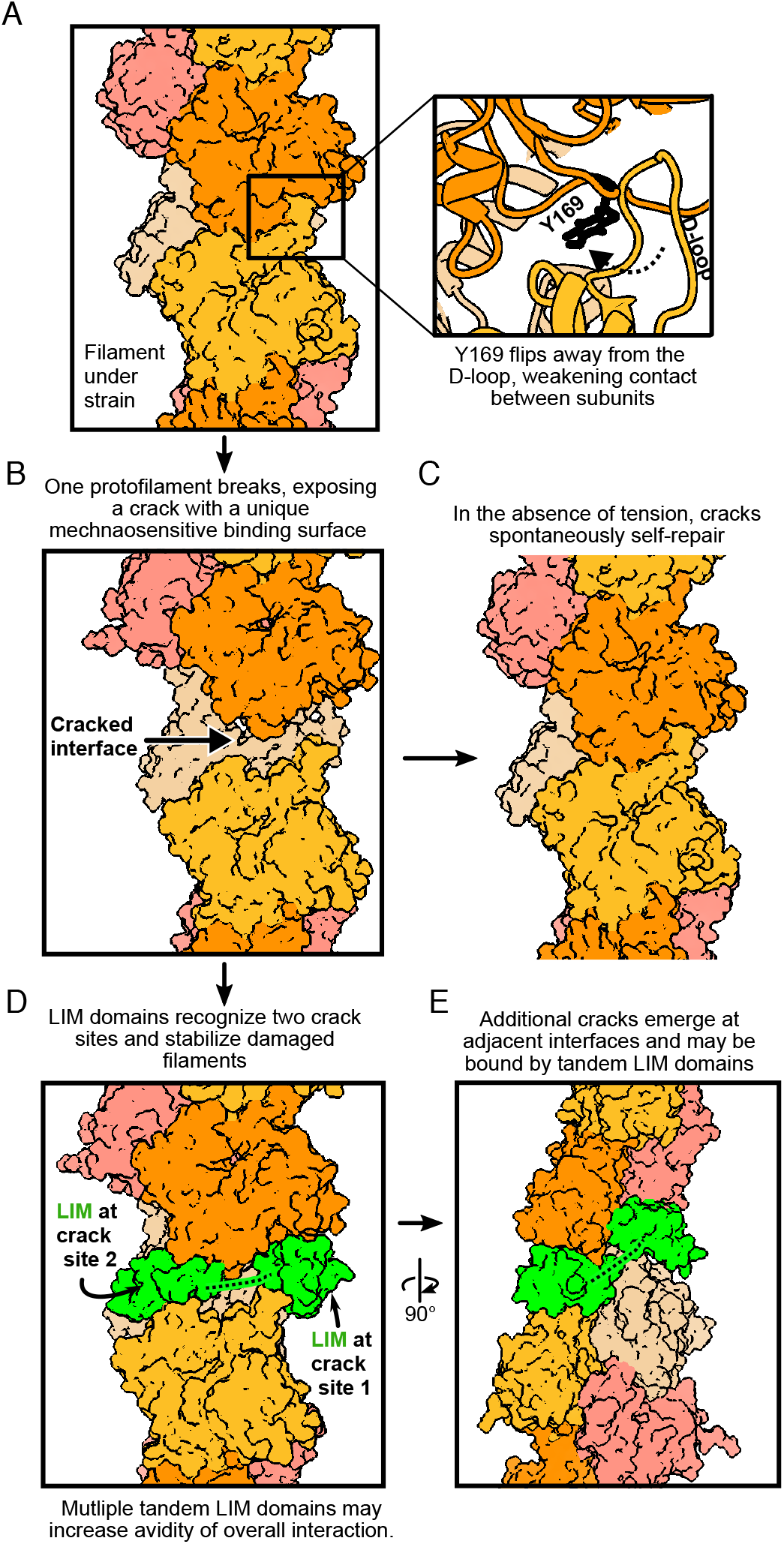
Cracked actin filaments as a mechanosensitive receptor. (A) Actin filaments subjected to mechanical strain break at interfaces where Y169 has flipped away from the D-loop of the neighboring subunit. Y169 flipping occurs more frequently at higher strains. (B) One protofilament breaks, while the other maintains all of its longitudinal connections, resulting in a cracked interface with a unique binding surface. (C) Cracks spontaneously self-repair when tension is no longer applied. (D) Cracks are mechanosensitive binding sites. LIM domains (green) can bind cracked interfaces in two non-mutually exclusive locations. Cracked interfaces with bound LIM domains withstand applied strain longer than without LIM bound. Dashed line represents the flexible linker between tandem LIM domains. (E) Under continued strain, additional cracks can form on the opposite protofilament strand. Tandem LIM domains may be able to bind cracks of two neighboring interfaces simultaneously, providing additional stability to the filament by binding both protofilaments. Dashed line represents the flexible linker between tandem LIM domains.

Given the importance of residue Y169, one could imagine that mutations away from tyrosine or phenylalanine result in deleterious effects. One experimental study explored the naturally-occurring zebrafish mutation Y169S (24). *In vivo*, the phenotype is a loss of filamentous actin. *In vitro*, the equivalent yeast actin mutant F169S initially nucleates and polymerizes, but rapidly depolymerizes into fragments thereafter. This behavior is rescued by introducing phalloidin, a molecule that stabilizes actin filaments. Our results suggest a mechanism to explain the surprising behavior of this mutation. If F169S actin initially forms a favorable connection between residue 169 and the D-loop of the adjacent subunit, it may be able to nucleate and polymerize. However, if the rate at which S169 loses its connection with the D-loop is much higher than WT (e.g., by flipping quickly under thermal fluctuations), then it will rapidly break apart into fragments. Phalloidin, by reinforcing connections between subunits, would stabilize the filament enough to overcome the unfavorable orientation of S169 and suppress the resulting fragmentation.

In the absence of applied strain, cracked actin filaments spontaneously repair themselves on short timescales without the need for chaperone proteins (Fig. 1D, 5C). This self-healing behavior is intuitive because the cracked interface consists of a barbed end and pointed end of neighboring subunits in close proximity and proper orientation with respect to each other due to the intact contacts made by the opposing protofilament. Given that barbed end elongation is diffusion limited, two nearby subunits oriented favorably, as seen at cracked interfaces, would be expected to establish stable inter-subunit contacts quickly in the absence of strain.

We have also simulated actin filaments in the ADP nucleotide state, as this represents the most aged filaments and likely matches the nucleotide state of the actin filaments in the *in vitro* experiments that inspired this work. It would be interesting to investigate whether the observations we have made here are modulated by the nucleotide state of actin. For example, if the prevalence that Y169 is in the flipped orientation is a function of ATP-hydrolysis and inorganic phosphate release, this likely in turn alters the prevalence of filament cracking for filaments in these nucleotide states. Given that ADP-actin filaments are less stable than ATP-actin filaments (25-27), yet structures of F-actin in all nucleotide states are nearly identical (28), perhaps differences in dynamics and prevalence of rare states, such as Y169’s flipping, can explain the observed differences in filament stability.

### Cracked interfaces are mechanosensitive receptors

Cracked interfaces present a unique force-induced binding surface (Fig. 3B, S3, 5B), while subunits at strained-intact interfaces appear remarkably similar to subunits in unstrained actin filaments (Fig. 1C, 3B). In particular, amino acids do not generally become more exposed prior to crack formation (Fig. 3C, S2). Therefore, we do not believe the strained-intact state is likely to contribute to force-activated binding of ABPs. More directly, our docking simulations included several strained-intact interfaces in addition to a cracked interface, and LIM domains were unlikely to bind them (Fig. 4B, S1). For these reasons, we propose that binding to cracked interfaces may be a general mechanism for mediating force-activated interactions between actin filaments and mechanosensitive ABPs. In particular, the hydrophobic and charged residues that become exposed in the D-loop (V43, M44, V45), W-loop (E167, Y169), and elsewhere (Q246, I287) drive the favorable interactions between LIM domains and the cracked interface in protein-protein docking simulations (Fig. 3, S3, S5, S6).

We further predict that resolving cracks in a frozen state using cryo-EM will be difficult. Cracks are rare, transient events. Given that cryo-EM depends on the averaging over many observations of roughly the same structure, collecting a large number of micrographs of cracked actin filaments could pose a significant challenge. Additionally, although the separation between subunits at cracked interfaces is noticeable in atomistic MD simulations, this separation at the EM micrograph level is likely to be very subtle, and identifying these occurrences will pose another challenge.

As noted earlier, a recent cryo-EM study reported the structure of bent actin filaments (12). At the subunit level, the changes with respect to unstrained filaments were subtler than the crack we observed in MD simulations. We believe that the structures solved by cryo-EM are most likely akin to the strained-intact state that we describe in this manuscript. This would be the much more common state for interfaces to be in in EM micrographs and displays much more subtle changes to the filament surface.

However, cracked interfaces with LIM domains present may prove easier to identify. For one, these structures are longer-lived than cracked interfaces with no LIM domains bound (Fig. 4E and F), so they will likely be better represented in the dataset. Additionally, the presence of bound LIM domains may aid in the identification of these structures in EM micrographs, especially if LIM domains bind to cracked interfaces at multiple neighboring interfaces and wrap around the filament (Fig. 5E).

### LIM domains recognize cracks

In docking simulations, we observed that an evolutionarily diverse library of 43 LIM domains associated with cracked interfaces over other regions of the filament surface, such as strained intact interfaces and filament ends (Fig. 4B and S1). This is consistent with experimental evidence that LIM domains can bind strained actin filaments (14-16). Furthermore, we observe two non-mutually exclusive binding sites at a single cracked interface (Fig. 4C, 5D), suggesting a mechanistic explanation for why three or more LIM domains in tandem separated by 7-8 amino acid linkers is necessary for mechanosensitivity of the vast majority of LIM domain proteins (15). First, the ∼ 4 nm separation between crack sites 1 and 2 is consistent with the separation between LIM domains in structures predicted by AlphaFold2, indicating a single LIM-containing protein could have multiple domains bound simultaneously at one cracked interface. Therefore, when one LIM domain associates with a crack site, unbound tandem LIM domains undergo tethered diffusion near a second LIM-binding crack site, boosting the avidity of the overall binding interaction (Fig. 5D). By tuning the linker lengths, the diffusive search is better constrained. Secondly, when LIM domains are bound to a cracked interface and the filament remains under tension, we find that crack site 2 can become exposed on the opposite protofilament strand prior to fragmentation (Fig. 4G). By having free LIM domains undergoing tethered diffusion, a single LIM-containing protein bound to the crack on one protofilament can quickly associate with the newly exposed crack on the opposite protofilament strand, allowing the LIM-containing protein to effectively wrap around the filament and reinforce lateral connections (Fig. 5E). As this process repeats, cracked interfaces stabilized by LIM domains may propagate along the filament, which may explain the spreading of LIM domains on actin filaments *in vitro* (14).

LIM domains bound to the crack substantially prolong the time before damaged actin filaments fragment (Fig. 4E and F). In addition to stabilizing individual filaments, this helps maintain intact actin filament networks in two ways. First, this prolongs the period that LIM domains can signal to repair proteins that the filament is damaged, increasing the likelihood that repair occurs prior to fragmentation. Second, physiological strains may be intermittent in nature, meaning if LIM domains stabilize filaments during periods of highest strain that otherwise would have resulted in fragmentation, then the filament may not be subjected to strains likely to lead to rupture again.

## METHODS

### MD simulations of the actin 13-mer

We used the 13-mer actin filament system in the ADP-state from *Zsolnay PNAS 2020* (17) with an updated Charmm36m forcefield (29). Briefly, we patterned the ADP-F-actin subunits from PDB 6DJO (9) according to the reported rise and twist values and included waters near the catalytic center from previously equilibrated simulations of actin filaments. We used the autosolvate and autoionize plugins in VMD (30) to construct a 100 mM KCl solvent box with a minimum of 2.2 nm of TIP3P water separating the protein from its periodic image. We then performed sequential energy minimizations using NAMD (31) with a 10 kcal mol^−1^A□^−2^ harmonic constraint on 1) everything except buffer; 2) everything except buffer and protein sidechains; 3) only the bound nucleotide and Mg^2+^; and 4) only Mg^2+^. The system underwent a 1-ns heating protocol bringing the temperature from 0 to 310 K under constraint selection 1). This was followed by a series of five 400-ps constrained equilibrations with the force constant on constraint selection 1) being halved each iteration (k = 10, 5, 2.5, 1.25, 0.625 kcal mol^−1^A□^−2^) followed by a sixth constrained equilibration of k = 0.1 kcal mol^−1^A□^−2^ lasting 1 ns. The system was equilibrated without constraints for 60 ns. Production runs were performed on Department of Defense High Performance Computing systems using GROMACS version 2019.1 (32) in the isothermal-isobaric (constant NPT) ensemble with v-rescale temperature coupling, Parinello-Rahman pressure coupling, and the leapfrog integrator. The particle mesh Ewald sum method was used to calculate electrostatic interactions with a cutoff of 1.2 nm. Equal and opposite forces were applied on the Cα center of mass of the three terminal subunits at the barbed and pointed ends via a constant force scheme using the GROMACS pull code. In total, 55 production run simulations were performed of actin 13-mers at varying tensions (Table S1). The direction of the force was along the axis of the filament throughout the duration of the simulation.

### Docking simulations

We used the ClusPro2.0 web server to perform the docking simulations (18, 23). Briefly, for each simulation run, a receptor and ligand structure are given as input. The ligand molecule is rotated 70,000 times, and each rotation is translated to fully explore the receptor, generating ∼10^9^ unique orientations of the ligand with respect to the receptor. Each combination of rotation and translation is scored based on an estimate of the binding energy. The 1,000 orientations with the best score are selected from the original ∼10^9^ and are grouped into clusters of binding poses with an RMSD relative to one another less than 9 Å. Clusters are ranked according to the number of the top 1,000 poses that they contain. This typically yields 15 to 30 binding poses with vastly different cluster sizes. We determined the location of each output pose on the filament (e.g., barbed end, pointed end, cracked interface, side) and weighted them according to the cluster size (Fig. S1A). We used the experimental result that mechanosensitive LIM domains do not appear enriched at filament ends to further filter the LIM docking output by accepting only poses ranked more highly than the highest ranked filament end-binding pose (Fig. S1B). This yielded a filtered set of binding poses for each of the 43 LIM domains tested (Fig. S1C). We also used the following PDB structures as non-LIM ligands: profilin: 2PAV; cofilin: 6UBY; α-catenin: 6UPV; metavinculin: 6UPW; Lifeact: 7AD9; CapZ: 7PDZ; actin: 1NWK (6, 33-36).

### MD simulations of actin 7-mer

To compare the stability of cracked actin filaments with and without bound LIM domains, we constructed an actin filament system consisting of 7 actin subunits with a cracked interface in the middle of the filament. For simulations with LIM domains present, we included the top pose of the first LIM domain of testin at both crack site 1 and crack site 2. Zn^2+^ ions were positioned by aligning the Alphafold2 structure to the solution NMR structure of the third LIM domain of FHL2 (PDB 2D8Z). Coordinating cysteines were in the deprotonated state. The minimization, heating, equilibration, and force-application protocol was the same as described for the actin 13-mer filament system. In total, 14 production run simulations were performed of cracked actin 7-mers: 8 in the absence of a bound LIM domain and 6 in the presence of the first LIM domain of testin bound to crack sites 1 and 2 (Table S1).

### Analysis of MD simulations and docking structures

We report a distance between longitudinal neighboring subunits, the subunit-subunit distance, or ss_dist_, which is the distance between the center of mass of Cα atoms of subunits i and i-2 (Fig. 1A). Interfaces were considered cracked when the 2-ns moving average of the ss_dist_ exceeded 6.3 nm. Filaments were considered fragmented when the second time derivative of the filament end-to-end distance (i.e., its acceleration) reached a maximum, indicating all connections had been severed. The end-to-end distance was calculated using the center of mass of Cα atoms of the three terminal subunits at each filament end. Accessible surface area (ASA) calculations were performed in VMD using a probe radius of 4 Å to estimate the filament surface relevant for protein-binding. MDAnalysis was used to construct contact maps (37). Visualizations were performed in VMD and Chimera (30, 38).

## AUTHOR CONTRIBUTIONS

V.Z., M.L.G., D.R.K., and G.A.V. designed research; V.Z. performed research and analyzed data; V.Z., M.L.G., D.R.K., and G.A.V. interpreted results and wrote the paper.

## ACKNOWLEDGEMENTS

This work was supported in part through the Department of Defense Army Research Office through Multidisciplinary Research Initiative Grant W911NF1410403 (V.Z., M.L.G., D.R.K, and G.A.V.) and in part by the National Institutes of Health through Awards R01GM143792 (M.L.G.), R01GM079265 (D.R.K), and R01GM063796 (G.A.V.). The authors acknowledge The University of Chicago Research Computing Center and the U.S. Department of Defense High Performance Computing Modernization Program for providing computational resources. We thank Dr. Cristian Suarez, Kashmeera Baboolall, Dr. Stefano Sala, and Prof. Patrick W. Oakes for helpful discussions.

