## Supplemental Information for "Cracked actin filaments as mechanosensitive receptors"

### SUPPLEMENTAL FIGURES

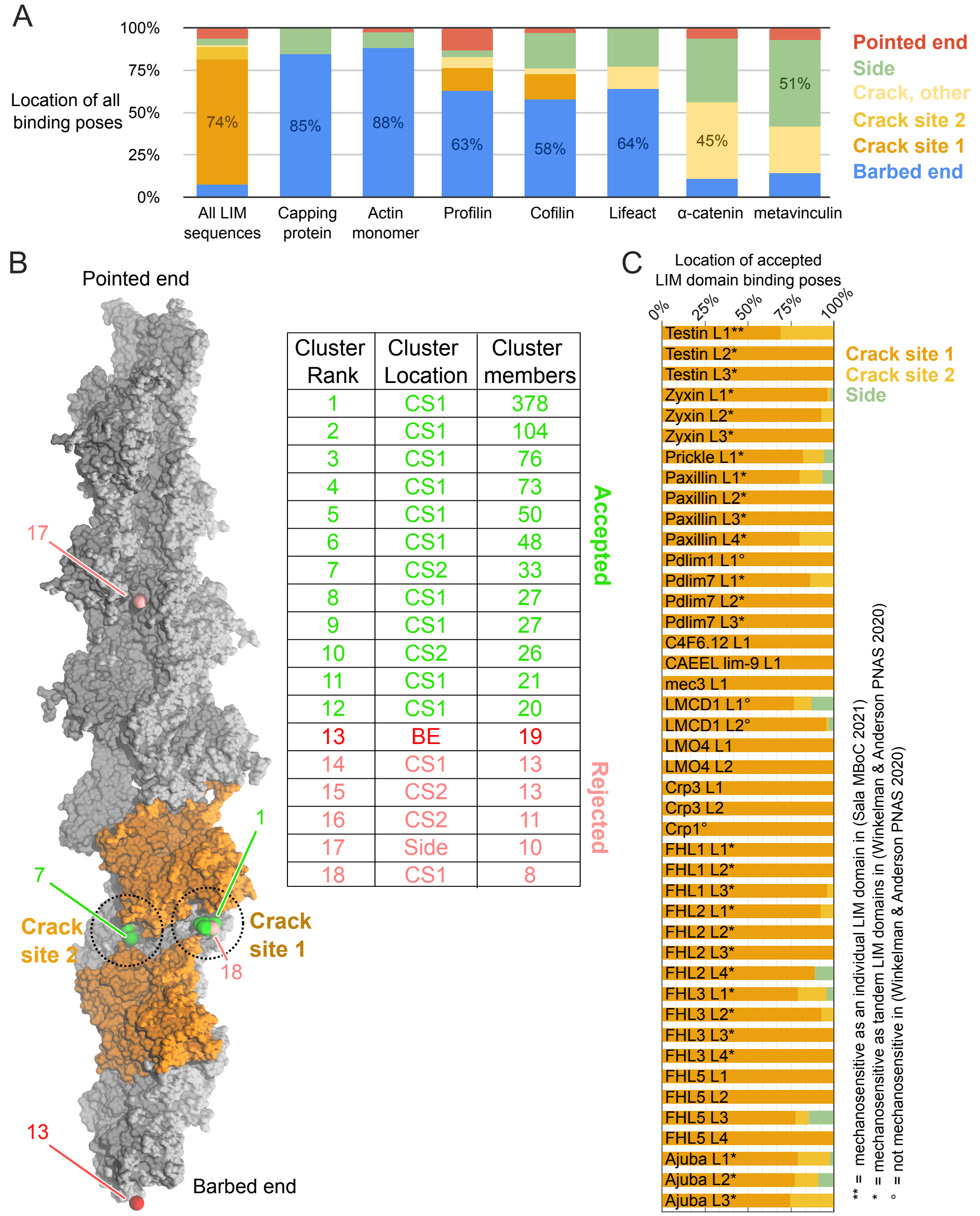


| Cluster Rank | Cluster Location | Cluster members |
|---|---|---|
| 1 | CS1 | 378 |
| 2 | CS1 | 104 |
| 3 | CS1 | 76 |
| 4 | CS1 | 73 |
| 5 | CS1 | 50 |
| 6 | CS1 | 48 |
| 7 | CS2 | 33 |
| 8 | CS1 | 27 |
| 9 | CS1 | 27 |
| 10 | CS2 | 26 |
| 11 | CS1 | 21 |
| 12 | CS1 | 20 |
| 13 | BE | 19 |
| 14 | CS1 | 13 |
| 15 | CS2 | 13 |
| 16 | CS2 | 11 |
| 17 | Side | 10 |
| 18 | CS1 | 8 |

Figure S1. Docking simulations of ABPs to a cracked actin filament.
(A) Location of all docking poses on a cracked actin 13-mer receptor. Each column corresponds to the indicated ligand. All 43 LIM domain sequences are shown at left as an aggregated dataset. The most likely binding location for each ligand is labeled as a percentage on the bar. Note that LIM domains have a striking preference for the cracked interface.
(B) Illustration of docking simulation processing. (Left) Space-filling model of the actin 13-mer receptor with subunits at the cracked interface highlighted in orange. Spheres represent the center of mass of each binding cluster output by ClusPro2.0 for a single LIM domain sequence. Several spheres are labeled with their ranking by cluster size. Color corresponds to acceptance/rejection.

(Right) Table of all binding poses. Poses with cluster size larger than the highest-ranked end-binding pose (red) are accepted (green) while others are rejected (pink). CS1/CS2 = crack site 1/2, BE = barbed end.
(C) Location of all accepted binding poses for each LIM domain sequence. In total, ~93% of accepted binding is at CS1, ~5% is at CS2, and <2% is on the filament side, not at the crack site.

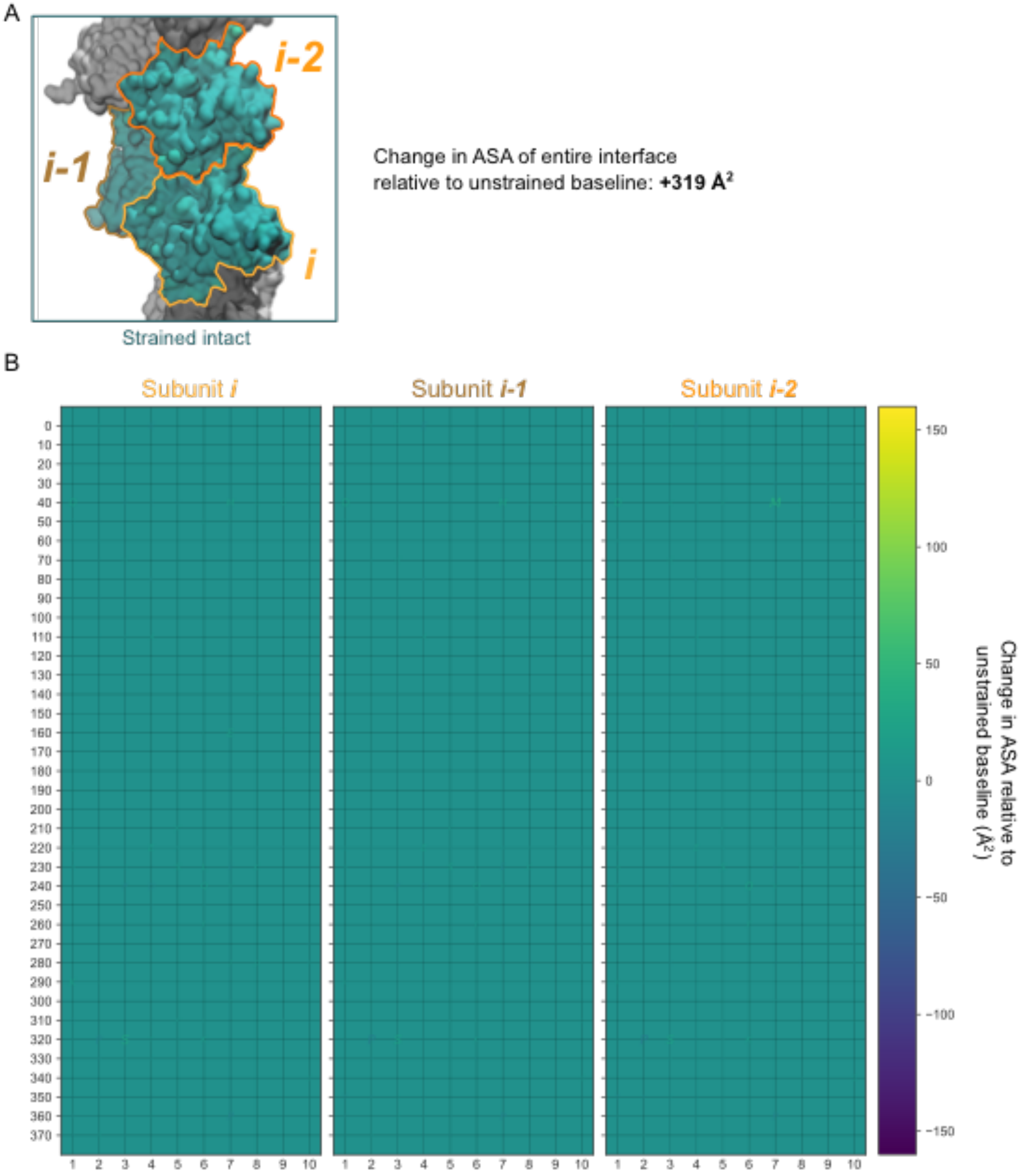


Figure S2. Strained-intact interfaces present a similar binding surface to interfaces in unstrained actin filaments.
(A) Space-filling representation of a representative strained-intact interface. Amino acid color corresponds to change in accessible surface area (ASA) relative to unstrained actin filaments according to the color bar in (B).
(B) Three grids list every amino acid in subunits i, i-1, and i-2 at a strained-intact interface. The size and color of each amino acid is scaled based on the change in ASA. Given that all residues

in strained-intact filaments maintain roughly the same ASA of unstrained filaments, these grids appear empty. The ASA probe radius is 4 Å. n = 22 simulations.

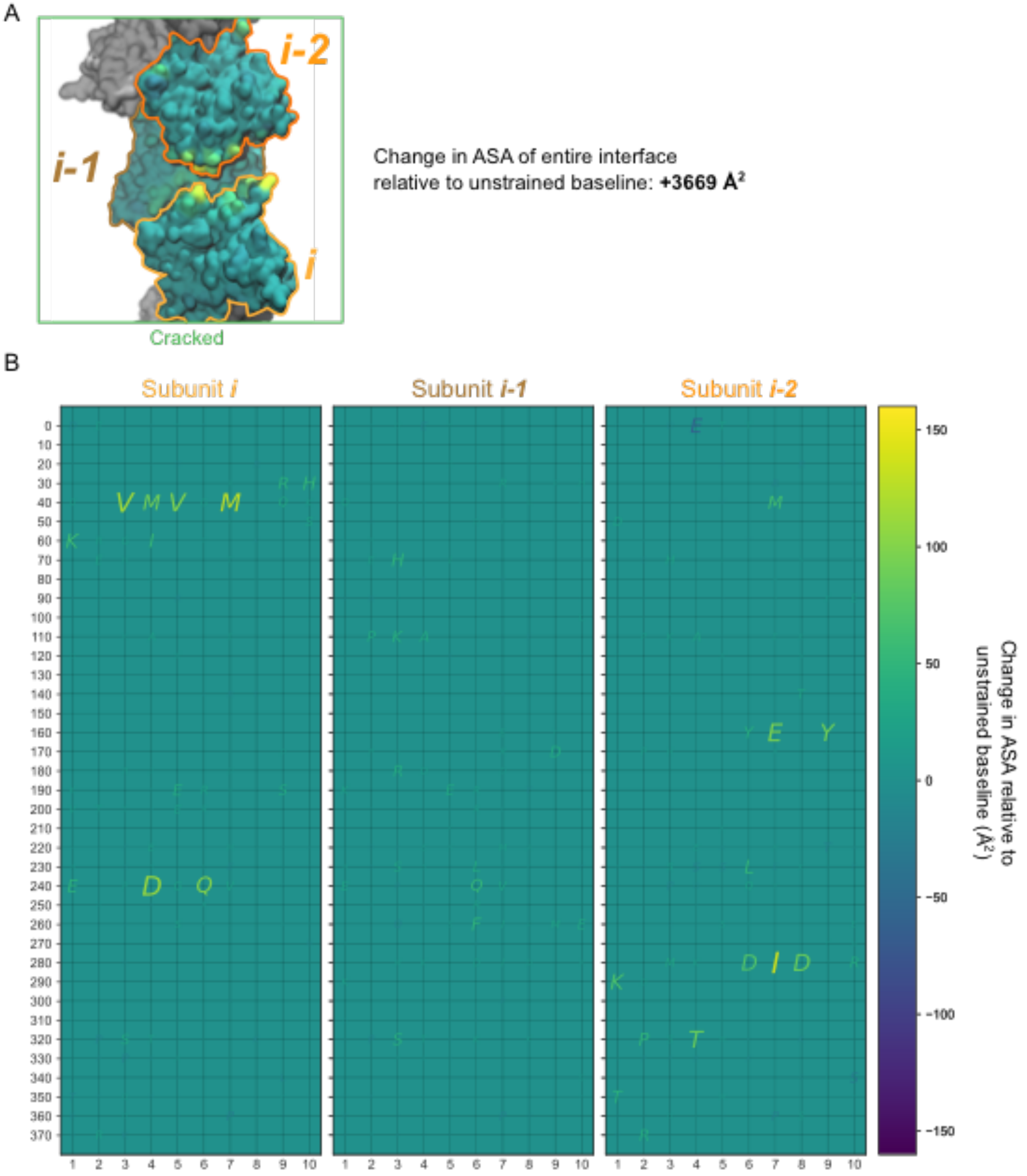


Figure S3. Cracked interfaces present a unique strain-induced binding surface.
(A) Space-filling representation of a representative cracked interface. Amino acid color corresponds to change in accessible surface area (ASA) relative to unstrained actin filaments according to the color bar in (B).
(B) Three grids list every amino acid in subunits i, i-1, and i-2 at a cracked interface. The size and color of each amino acid is scaled based on the change in ASA. The ASA probe radius is 4 Å. n = 45 simulations.

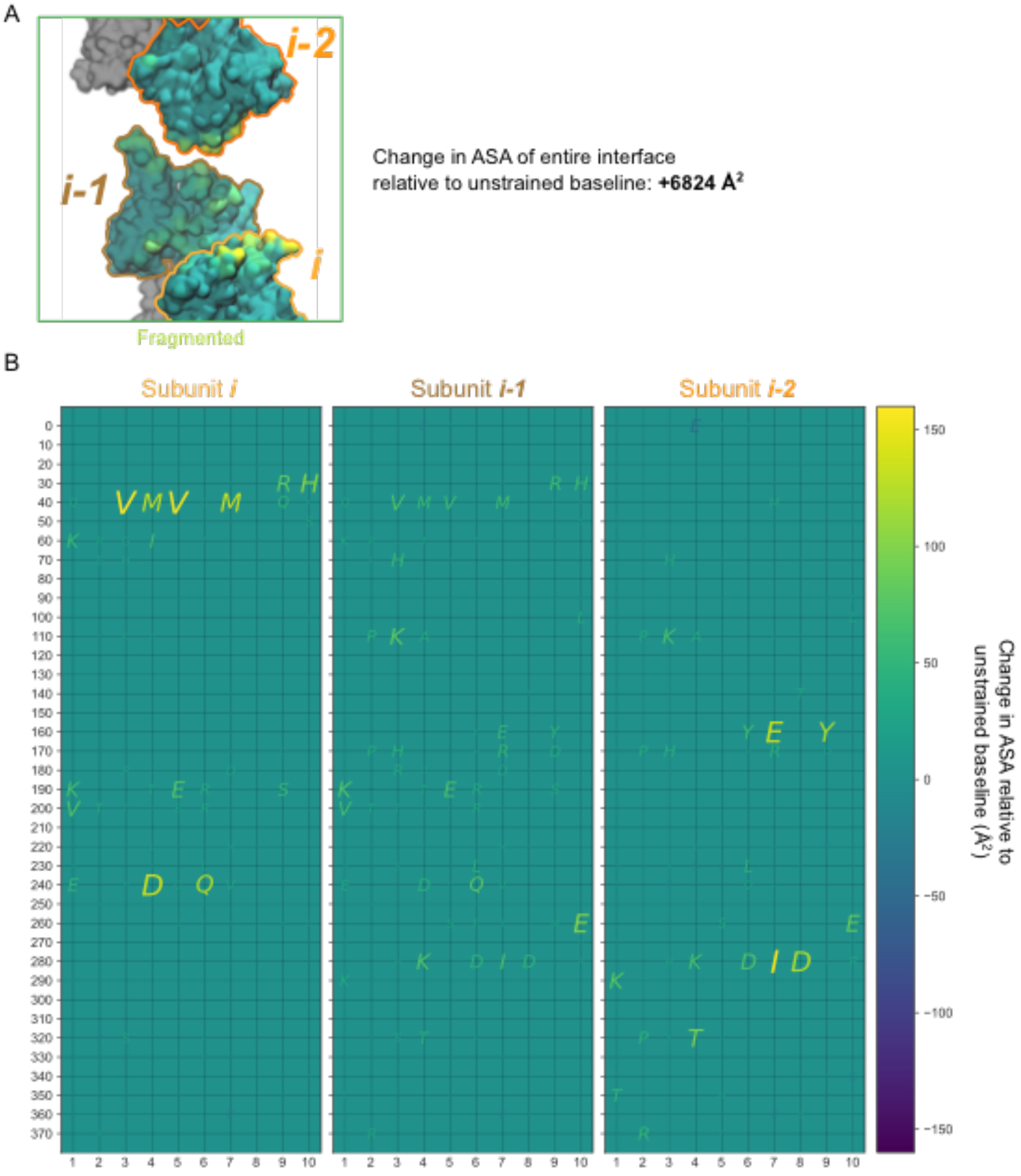


Figure S4. Filament fragmentation exposes typically-buried amino acids.
(A) Space-filling representation of a representative cracked interface. Amino acid color corresponds to change in accessible surface area (ASA) relative to unstrained actin filaments according to the color bar in (B).
(B) Three grids list every amino acid in subunits i, i-1, and i-2 at a fragmented interface. The size and color of each amino acid is scaled based on the change in ASA. The ASA probe radius is 4 Å. n = 45 simulations.

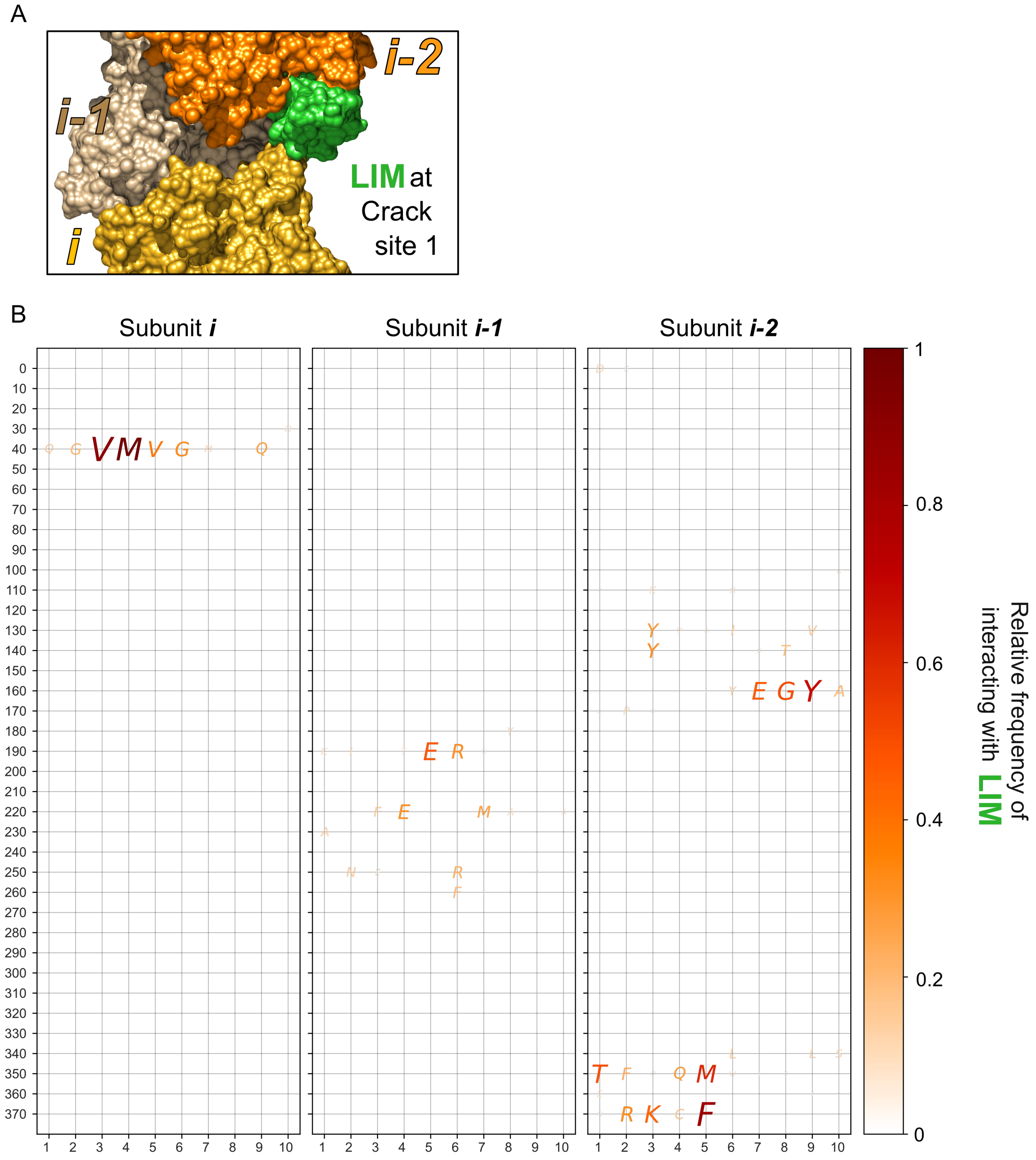


Figure S5. LIM domains bound to crack site 1 interact with residues that become exposed at cracked interfaces.

(A) Space-filling representation of a LIM domain (green) bound to the cracked interface at crack site 1.

(B) Three grids list every amino acid in subunits i, i-1, and i-2 at the cracked interface. The size and color of each amino acid is scaled based on its relative frequency of interacting with LIM bound to crack site 1 across all accepted binding poses of all 43 LIM domain sequences. Contact was calculated using a 5 Å cutoff.

A

B

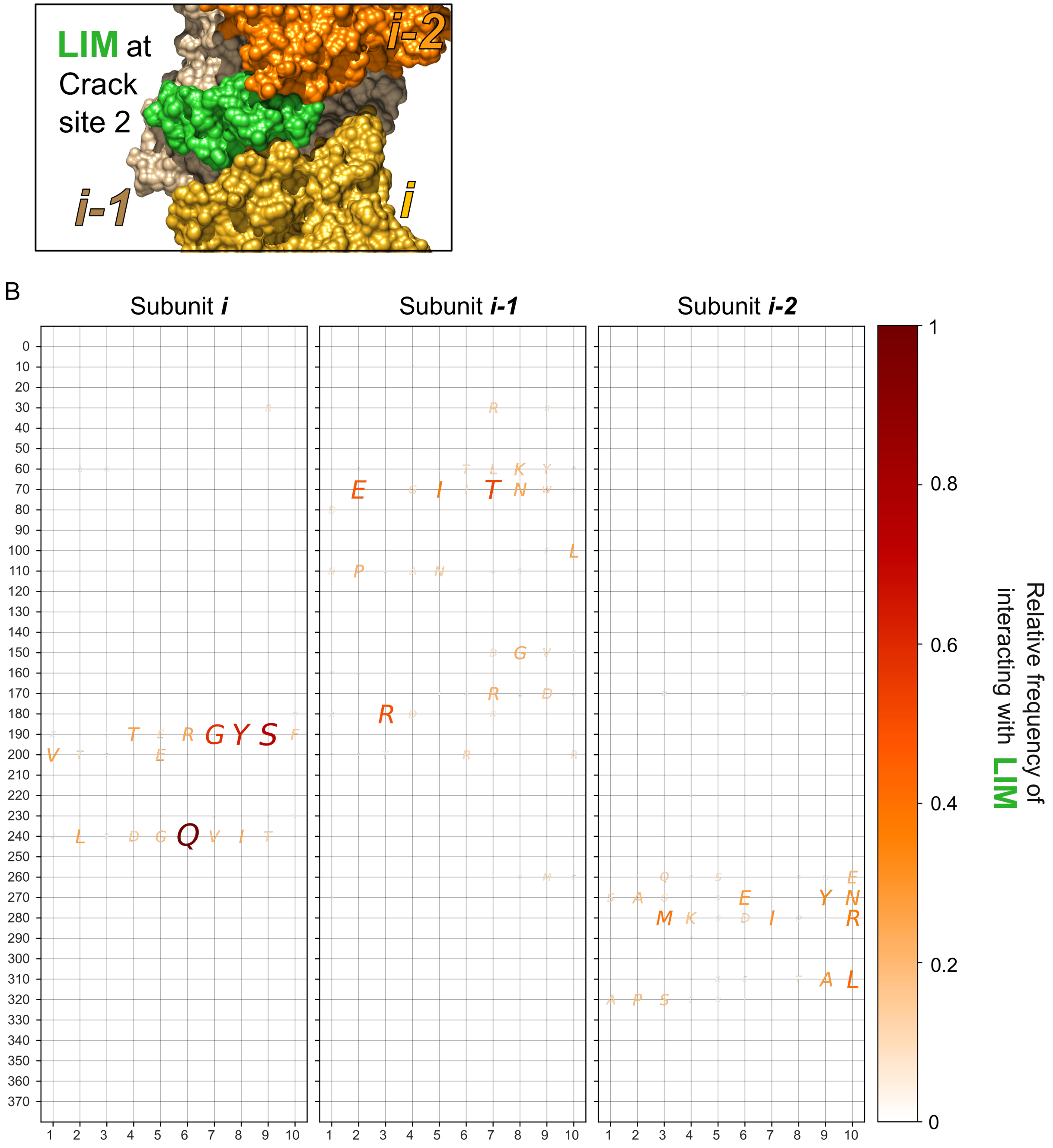


Figure S6. LIM domains bound to crack site 2 interact with residues that become exposed at cracked interfaces.

(A) Space-filling representation of a LIM domain (green) bound to the cracked interface at crack site 2.

(B) Three grids list every amino acid in subunits i, i-1, and i-2 at the cracked interface. The size and color of each amino acid is scaled based on its relative frequency of interacting with LIM bound to crack site 2 across all accepted binding poses of all 43 LIM domain sequences. Contact was calculated using a 5 Å cutoff.

Movie S1. Actin filaments crack before they fragment. An actin filament under 500 pN tension initially maintains all connections between subunits. Eventually a longitudinal interface on one protofilament loses contact, resulting in a metastable cracked conformation. Under continued strain, the opposite protofilament breaks and the filament fully fragments. The duration is 123 ns and frames are smoothed using a 3-ns window to clarify dominant motions.

Movie S2. The flip of Y169 (black) of subunit i-2 (yellow) away from the D-loop of subunit i (orange) precedes the first longitudinal interface breaking (crack formation). Frames are smoothed using a 3-ns window to clarify dominant motions. The duration is 100 ns, and the filament is subjected to a 550 pN tension.

Movie S3. Crack site 2 becomes exposed before fragmentation of an already-cracked filament stabilized by LIM domains. The last 170 ns are shown of a 2.43 μs simulation of an actin filament with two LIM domains (green) bound to crack site 1 and 2 at the interface of subunits i (yellow) and i-2 (orange). Before fragmentation occurs, crack site 2 becomes exposed on the neighboring interface of the opposite protofilament strand between subunits i-1 (tan) and i-3 (pink, *top*). This may allow an unbound tandem LIM domain (not shown) to bind cracks on both protofilament strands simultaneously. Frames are smoothed using a 1-ns window to clarify dominant motions. The filament is subjected to 250 pN tension.

**SUPPLEMENTAL TABLE**

Table S1: Simulations performed in this study

| Simulation number | Initial structure | Number of actin subunits | Tension (pN) | Replicate number | Duration (ns) | Final state |
|---|---|---|---|---|---|---|
| 1 | 6DJO | 13 | 0 | 1 | 373 | Intact |
| 2 | 6DJO | 13 | 0 | 2 | 988 | Intact |
| 3 | 6DJO | 13 | 0 | 3 | 580 | Intact |
| 4 | 6DJO | 13 | 200 | 1 | 1,091 | Intact |
| 5 | 6DJO | 13 | 200 | 2 | 300 | Intact |
| 6 | 6DJO | 13 | 400 | 1 | 810 | Intact |
| 7 | 6DJO | 13 | 400 | 2 | 400 | Intact |
| 8 | 6DJO | 13 | 500 | 1 | 91 | Fragment |
| 9 | 6DJO | 13 | 500 | 2 | 123 | Fragment |
| 10 | 6DJO | 13 | 500 | 3 | 340 | Fragment |
| 11 | 6DJO | 13 | 500 | 4 | 307 | Fragment |
| 12 | 6DJO | 13 | 500 | 5 | 251 | Fragment |

| | | | | | | |
|---|---|---|---|---|---|---|
| 13 | 6DJO | 13 | 500 | 6 | 510 | Fragment |
| 14 | 6DJO | 13 | 500 | 7 | 180 | Fragment |
| 15 | 6DJO | 13 | 500 | 8 | 110 | Fragment |
| 16 | 6DJO | 13 | 550 | 1 | 200 | Fragment |
| 17 | 6DJO | 13 | 550 | 2 | 100 | Fragment |
| 18 | 6DJO | 13 | 550 | 3 | 80 | Fragment |
| 19 | 6DJO | 13 | 550 | 4 | 194 | Fragment |
| 20 | 6DJO | 13 | 550 | 5 | 72 | Fragment |
| 21 | 6DJO | 13 | 600 | 1 | 40 | Fragment |
| 22 | 6DJO | 13 | 600 | 2 | 82 | Fragment |
| 23 | 6DJO | 13 | 600 | 3 | 87 | Fragment |
| 24 | 6DJO | 13 | 600 | 4 | 30 | Fragment |
| 25 | 6DJO | 13 | 600 | 5 | 40 | Fragment |
| 26 | Cracked | 13 | 0 | 1 | 57 | Intact |
| 27 | Cracked | 13 | 0 | 2 | 142 | Intact |
| 28 | Cracked | 13 | 0 | 3 | 101 | Intact |
| 29 | Cracked | 13 | 300 | 1 | 171 | Fragment |
| 30 | Cracked | 13 | 300 | 2 | 141 | Fragment |
| 31 | Cracked | 13 | 300 | 3 | 55 | Fragment |
| 32 | Cracked | 13 | 300 | 4 | 358 | Fragment |
| 33 | Cracked | 13 | 300 | 5 | 60 | Fragment |
| 34 | Cracked | 13 | 350 | 1 | 30 | Fragment |
| 35 | Cracked | 13 | 350 | 2 | 70 | Fragment |
| 36 | Cracked | 13 | 350 | 3 | 70 | Fragment |
| 37 | Cracked | 13 | 350 | 4 | 100 | Fragment |
| 38 | Cracked | 13 | 400 | 1 | 40 | Fragment |
| 39 | Cracked | 13 | 400 | 2 | 20 | Fragment |
| 40 | Cracked | 13 | 400 | 3 | 27 | Fragment |
| 41 | Cracked | 13 | 400 | 4 | 40 | Fragment |
| 42 | Cracked | 13 | 400 | 5 | 40 | Fragment |
| 43 | Cracked | 13 | 450 | 1 | 20 | Fragment |
| 44 | Cracked | 13 | 450 | 2 | 30 | Fragment |
| 45 | Cracked | 13 | 450 | 3 | 20 | Fragment |

| | | | | | | |
|---|---|---|---|---|---|---|
| 47 | Cracked | 13 | 500 | 1 | 30 | Fragment |
| 47 | Cracked | 13 | 500 | 2 | 18 | Fragment |
| 48 | Cracked | 13 | 550 | 1 | 20 | Fragment |
| 49 | Cracked | 13 | 550 | 2 | 20 | Fragment |
| 50 | Cracked | 13 | 550 | 3 | 20 | Fragment |
| 51 | Cracked | 13 | 600 | 1 | 11 | Fragment |
| 52 | Cracked | 13 | 600 | 2 | 10 | Fragment |
| 53 | Cracked | 13 | 600 | 3 | 10 | Fragment |
| 54 | Strained-intact | 13 | 500 | 1 | 64 | Fragment |
| 55 | Strained-intact | 13 | 550 | 1 | 30 | Fragment |
| 56 | Cracked | 7 | 250 | 1 | 70 | Fragment |
| 57 | Cracked | 7 | 250 | 2 | 20 | Fragment |
| 58 | Cracked | 7 | 250 | 3 | 50 | Fragment |
| 59 | Cracked | 7 | 250 | 4 | 50 | Fragment |
| 60 | Cracked | 7 | 250 | 5 | 50 | Fragment |
| 61 | Cracked | 7 | 500 | 1 | 11 | Fragment |
| 62 | Cracked | 7 | 500 | 2 | 10 | Fragment |
| 63 | Cracked | 7 | 500 | 3 | 5 | Fragment |
| 64 | Cracked+LIM[1] | 7 | 250 | 1 | 1,479 | Fragment |
| 65 | Cracked+LIM | 7 | 250 | 2 | 2,430 | Fragment |
| 66 | Cracked+LIM | 7 | 250 | 3 | 370 | Fragment |
| 67 | Cracked+LIM | 7 | 500 | 1 | 21 | Fragment |
| 68 | Cracked+LIM | 7 | 500 | 2 | 15 | Fragment |
| 69 | Cracked+LIM | 7 | 500 | 3 | 15 | Fragment |
| **Total** | — | — | — | — | **13,800** | — |

1. System composed of a cracked actin 7-mer with two copies of the first LIM domain of testin bound to crack sites 1 and 2.
